# Glucosylceramide synthase is required for C6-ceramide nanoliposome-induced organelle stress and cell death

**DOI:** 10.64898/2026.09.21.753214

**Authors:** Luke R. Vass, Ella W. Smith, Jayda M. Cavanaugh, Summer L. Johnson, Ariana Sabzevari, Pedro Costa-Pinheiro, Jeremy J.P. Shaw, Thomas P. Loughran, Todd E. Fox

## Abstract

Sphingolipids are bioactive lipids that regulate key signaling pathways both directly as ligands and through membrane re-organization. Ceramide sits at the center of this network and is considered pro-death in many contexts, making ceramide accumulation an attractive therapeutic strategy. Given the wide-ranging regulation of cellular responses this network exerts, better understanding ceramide metabolism may promote therapeutic efficacy of sphingolipid-based therapeutics. Ceramide glycosylation, catalyzed by glucosylceramide synthase (GCS; UGCG), is in turn widely regarded as a detoxification route thus limiting efficacy of ceramide-based therapeutics. Here we show the opposite. Delivery of short-chain C6-ceramide via a ceramide nanoliposome (CNL) induced organelle stress and cell death in chronic lymphocytic leukemia (CLL) via the accumulation of glycosphingolipids (GSLs), rather than through ceramide itself. Pharmacologic and genetic blockade of GCS protected B-cell leukemia, breast carcinoma, glioblastoma, lung adenocarcinoma, and non-malignant embryonic kidney cells from CNL-induced death. Conversely, exogenous C8-glucosylceramide was sufficient to kill cells that cannot degrade it. We show that GSL accumulation drives an ordered organelle response beginning with lysosomal deacidification, endoplasmic reticulum stress, followed by mitochondrial respiratory capacity decline, each attenuated by inhibition of GSL synthesis. These effects were accompanied by MLKL phosphorylation, increased activity of the stress sensor JNK and CHOP induction, with JNK inhibition partially protecting from death. These findings invert the prevailing view of ceramide glycosylation as a resistance mechanism and identify glycosphingolipid flux as a required effector arm of ceramide-directed therapy.

## Introduction

Chronic lymphocytic leukemia (CLL) is one of the most commonly diagnosed adult leukemias in Western countries and is characterized by the accumulation of malignant B cells, leading to cytopenias, lymphadenopathy, and progressive immune dysfunction.^1^ Frontline therapy is now centered on two major mechanisms: continuous inhibition of B-cell receptor (BCR) signaling with Bruton tyrosine kinase (BTK) inhibitors and inhibition of the anti-apoptotic protein BCL2 with venetoclax.^2–4^ These approaches have largely displaced chemoimmunotherapy and provide durable disease control for many patients. However, patients whose disease progresses after exposure to both a covalent BTK inhibitor and venetoclax, often termed double-refractory CLL, have limited treatment options and a median overall survival of approximately two years.^5^ Resistance can arise through BTK mutations that preserve scaffolding function and through BCL2 mutations that alter the BH3-binding groove.^6^ Therapeutic strategies that induce cell death independently of BCR signaling and BCL2 are therefore of interest.

Sphingolipid metabolism provides one such potential route. Ceramide is a central intermediate in sphingolipid metabolism and can promote cell death in diverse cellular contexts. Ceramide nanoliposomes (CNL), a biocompatible delivery system for short-chain C6-ceramide, have demonstrated activity across preclinical cancer models,^7,8^ completed a phase I study in advanced solid tumors,^9^ and are currently undergoing clinical evaluation in hematologic malignancies (NCT04716452). We have previously shown that CNL induces cell death in CLL cells and limits tumor progression in a CLL xenograft model.^8,10^

CNL, however, does not deliver ceramide into a metabolically static environment. Following cellular uptake, C6-ceramide can undergo phosphorylation, glycosylation, conversion to sphingomyelin, and deacylation and reacylation of the sphingoid backbone, generating metabolites with distinct biological activities.^11^ The sphingolipid metabolic state of the target cell also influences its response to CNL.^12^ Thus, the effects of CNL may depend not only on delivery of the ceramide itself but also on its subsequent metabolic fate. Importantly, the contribution of these metabolic pathways to CNL-induced cell death has not been established.

Glycosylation is particularly relevant because glucosylceramide synthase (GCS; *UGCG*) initiates conversion of ceramide into glucosylceramide and downstream complex glycosphingolipids (GSLs). A common model of ceramide biology predicts that this pathway diminishes ceramide-mediated cytotoxicity by diverting ceramide into less toxic metabolites, and *UGCG* inhibition has consequently been investigated as a strategy to enhance the activity of anticancer therapies.^13,14^ Whether this model applies to direct therapeutic delivery of ceramide is unknown. However, GSLs are biologically active lipids that participate in membrane organization and signaling,^15^ raising the possibility that their synthesis could contribute to, rather than oppose, the cellular response to CNL. Sustained GSL accumulation also disrupts organelle function, as seen in glycosphingolipid storage disorders, where progressive accumulation produces lysosomal, endoplasmic reticulum, and mitochondrial dysfunction.^16,17^ This distinction is particularly relevant in CLL, where *UGCG*-dependent sphingolipid metabolism has been linked to BCR signaling, drug sensitivity, and clinical outcome.^18,19^

Here, we tested whether GSL synthesis protects CLL cells from CNL or contributes to its cytotoxic activity. We find that GSL synthesis is required for CNL-induced organelle stress and cell death. This requirement extends across multiple malignant and non-malignant cell types, indicating that metabolism of delivered ceramide into GSLs is not a route of drug inactivation, but rather an essential therapeutic component of ceramide delivery.

## Results

### Ceramide nanoliposome (CNL)-induced cell death follows the conversion of ceramide into glycosphingolipids

We previously showed that CNL-delivered C6-ceramide (**Fig. 1A, insets 1 and 2**) induces cell death in CLL cells and limits tumor progression *in vivo*.^8,10^ Consistent with this, CNL reduced JVM-3 viability in a concentration- and time-dependent manner **(Fig. 1B)**. Intracellular ATP was progressively depleted over the same period, with small reductions detectable within the first hours of treatment and a loss of approximately half by 22 hours **(Fig. 1C)**. To define the metabolic events associated with this response, we asked how delivered C6-ceramide is metabolized by the sphingolipid network **(Fig. 1A)** across the same timeframe.

**Figure 1:**
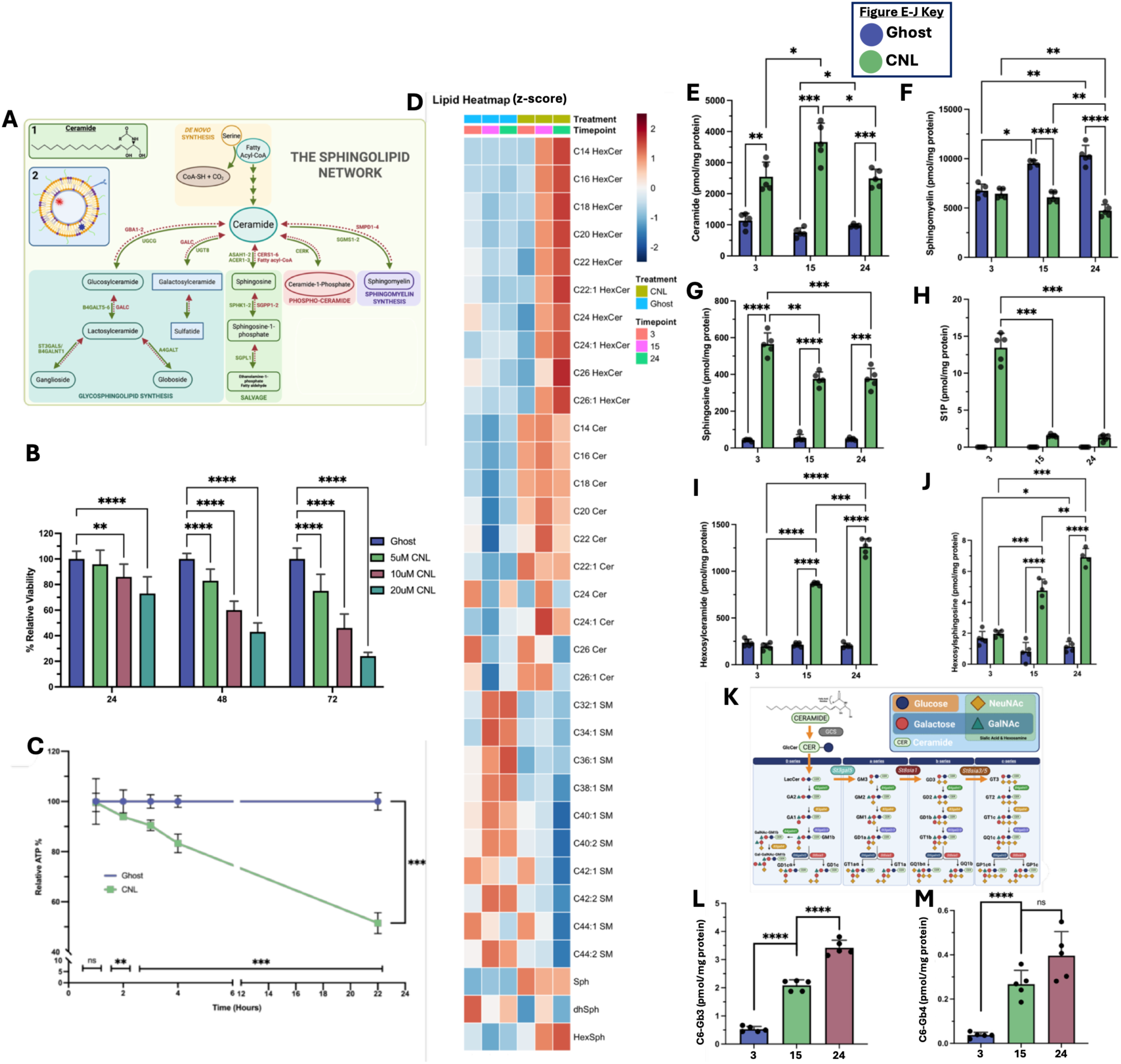
Ceramide nanoliposomes (CNL) induce time-dependent loss of viability and remodeling of the sphingolipid network. **(A)** Schematic of key sphingolipid metabolic pathways, with insets showing the structure of ceramide **(1)** and a representative nanoliposome **(2)**. **(B)** MTS measured viability of JVM-3 cells treated with the indicated concentration of CNL, or 20 µM ghost (control) liposomes, for 24, 48 or 72 h. **(C)** Relative ATP levels in viable JVM-3 cells following treatment with 20 µM ghost liposomes or CNL, assayed by CellTiter-Glo. **(D)** Summary of LC-MS/MS lipidomic profiling of JVM-3 cells treated with 20 µM ghost or CNL for the indicated time; heatmap shows Z-scores of lipid quantifications. **(E–J)** LC-MS/MS quantification of ceramides **(E)**, sphingomyelins **(F)**, sphingosine **(G)**, sphingosine-1-phosphate (S1P, **H**), hexosylceramides **(I)** and hexosylsphingosines **(J)** following 20 µM CNL (green) or control liposomes (blue) at the indicated time. **(K)** Summary of GSL metabolites. **(L, M)** LC-MS/MS profiling of C6-Gb3 (L) and C6-Gb4 (M) over time following treatment with 20 µM CNL. Statistics: two-way ANOVA with Fisher’s LSD test (E–J) or Tukey’s post-hoc test (B); one-way ANOVA with Tukey’s post-hoc test (L, M); Student’s t-test (C). Each timepoint was normalized to its own control for (C). Error bars represent standard deviation of 3 (B, C) or 5 (E–M) biological replicates. *p<0.05, **p<0.01, ***p<0.001, ****p<0.0001.

Lipidomic profiling of CNL-treated cells by liquid chromatography–tandem mass spectrometry (LC-MS/MS) at 3, 15 and 24 hours **(summarized in Fig. 1D)** showed that total ceramides rose sharply by 3 hours, consistent with rapid uptake of delivered C6-ceramide, peaked at 15 hours and declined thereafter **(Fig. 1E)**. Sphingomyelins were likewise elevated at 3 and 15 hours before falling below vehicle by 24 hours **(Fig. 1F)**, while sphingosine **(Fig. 1G)** and sphingosine-1-phosphate **(Fig. 1H)** were elevated early and declined at later times, indicating that the sphingomyelin, salvage and phosphorylation arms are all engaged early with ceramide delivery, resolving by the time viability is lost. In contrast, hexosylceramide **(Fig. 1I)** and hexosylsphingosine **(Fig. 1J)** increased significantly over the same period and were still rising at 24 hours. Among the routes resolved by our panel, glycosylation was therefore the only one whose products accumulated over the interval in which viability and ATP were lost, rather than peaking before it.

Hexosylceramide, as measured here, includes both glucosylceramide and galactosylceramide. These lipids initiate divergent branches of GSL metabolism, with *UGCG* and galactosylceramide synthase (*UGT8*) as the respective entry enzymes. Glucosylceramide is the more abundant of the two and serves as the precursor for more complex GSLs, including gangliosides and globosides **(Fig. 1K)**. We therefore extended profiling to downstream species and found that CNL increased multiple GSLs through the pathway, including the C6 forms of the globosides Gb3 **(Fig. 1L)** and Gb4 **(Fig. 1M)**, which are not present in vehicle-treated cells and derive from the delivered C6-ceramide backbone.

### Glucosylceramide synthase activity is required for CNL-induced death

To test whether GSL synthesis contributes to CNL-induced death, we utilized ibiglustat (venglustat, GZ/SAR402671) a clinical-stage GCS inhibitor (**Fig. 2A**). Ibiglustat reduced basal hexosylceramide levels and prevented its accumulation following CNL treatment (**Fig 2B**), confirming target engagement and indicating that this pool is predominantly glucosylceramide, since ibiglustat inhibits GCS but not galactosylceramide synthase. Accumulation of downstream species C6-Gb3 (**Fig 2C**), C6-Gb4 (**Fig 2D**) and C6-GM3 (**Fig 2E**) was also nearly completely ablated.

**Figure 2:**
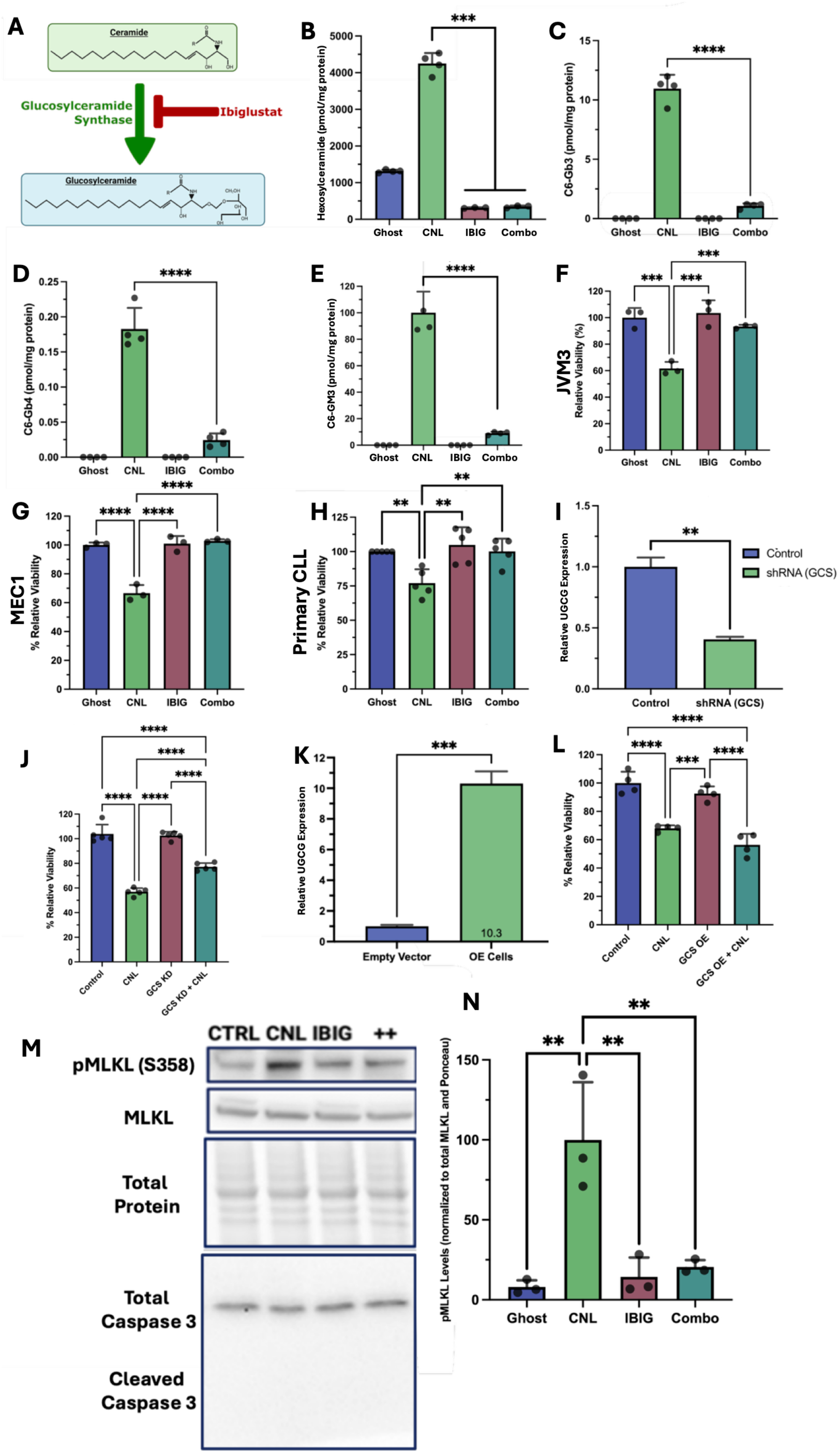
GSL accumulation is required for CNL-induced death. **(A)** Schematic of GCS activity and its pharmacological inhibition by ibiglustat. **(B–E)** LC-MS/MS quantification of hexosylceramide **(B)**, C6-Gb3 **(C)**, C6-Gb4 **(D)** and C6-GM3 **(E)**. **(F–H)** alamarBlue viability of JVM-3 **(F)**, MEC-1 **(G)** and primary CLL patient samples **(H)** treated with 20 µM ghost or CNL for 24 h, with or without a 1 h pre-treatment with 2 µM ibiglustat (IBIG). **(I, K)** *UGCG* expression by RT-qPCR following lentiviral transduction of shRNA (I) or overexpression **(K)** constructs into JVM-3 cells. **(J, L)** Viability of *UGCG*-knockdown **(J)** or *UGCG*-overexpressing **(L)** lines relative to vector-control JVM-3 cells in response to 20 µM CNL, assayed by alamarBlue. (M) Western blot for the indicated targets following the treatments above. **(N)** Quantification of MLKL phosphorylation, normalized to total MLKL and Ponceau. Statistics: one-way ANOVA with Tukey’s post-hoc test throughout. Lines expressing shRNA or the overexpression construct were normalized to their own vehicle control, so those comparisons are within-line rather than a test of a difference between lines. Error bars represent standard deviation; points represent biological replicates (n = 3 for F–H, J, L; n = 3 for I, K). *p<0.05, **p<0.01, ***p<0.001, ****p<0.0001.

Contrary to the expectation that suppressing this pathway would increase CNL efficacy, ibiglustat rescued both JVM-3 (**Fig 2F**) and MEC-1 (**Fig 2G**) CLL cell lines from CNL-induced death, restoring viability to near control levels. The same protection was observed in primary CLL patient samples (**Fig 2H**). Ibiglustat alone did not affect viability in any of the three CLL systems over this timeframe. To further confirm that this reflects a loss of GSL synthesis, we targeted *UGCG* genetically by shRNA. Partial knockdown (∼59.5%, **Fig. 2I)** recovered approximately 44% of the CNL-induced viability loss **(Fig. 2J)**, correlating with pharmacologic inhibition. Conversely, *UGCG* overexpression **(Fig. 2K)** did not increase sensitivity to CNL **(Fig. 2L)**.

GSL synthesis inhibition served to increase ceramide levels further (∼60%, **Fig S1A**) indicating that the toxicity of CNL is not driven by ceramide accumulation itself. Ceramide and death therefore move in opposite directions within a single experiment, which argues directly against ceramide accumulation as the proximate death signal. Sphingosine also increased moderately under GSL synthesis inhibition (**Fig S1B**), while sphingomyelin (**Fig S1C**), which showed depletion by CNL, was restored towards baseline.

To determine which cell death program GSL accumulation engages, we profiled cell death markers via western blot in JVM-3 cells. Phosphorylation of MLKL at S358, increased with CNL and returned to baseline with GSL synthesis inhibition (**Fig 2M, N**) while cleaved caspase 3 was not detected. MLKL phosphorylation is a marker of necroptotic signaling rather than a demonstration of it, and we expand on the question of death modality below.

### CNL induces a late transcriptional program of immune activation and cell stress

To define the cellular response to GSL accumulation, we profiled JVM-3 cells treated with CNL for 6 or 24 hours by bulk RNA-sequencing. Differential expression was limited and functionally incoherent at 6 h and converged on a coordinated stress and inflammatory program at 24 h, coinciding with the interval over which GSLs accumulate. The 24 h response was dominated by transcripts related to cell stress (*EGR1*, *ASS1*, *DDIT3*/CHOP, CHAC1) and immune activation (*CCL3*, *CCL4*, *IL23A*) (**Fig. 3A**), and over-representation analysis of the upregulated gene set was led by TNFα/NFκB and inflammatory response signatures (**Fig. 3B**). Footprint-based inference across the two timepoints placed the same pathways (**Fig. 3C**) and their associated transcription factors (**Fig. 3D**) among those most differentially active at 24 h. We confirmed induction of these transcripts by RT-qPCR and found that each was reduced when GSL synthesis was inhibited with ibiglustat (**Fig. 3E**), placing the inflammatory program downstream of GSL accumulation. Investigations into whether this program contributes to death are ongoing and not reported here.

**Figure 3:**
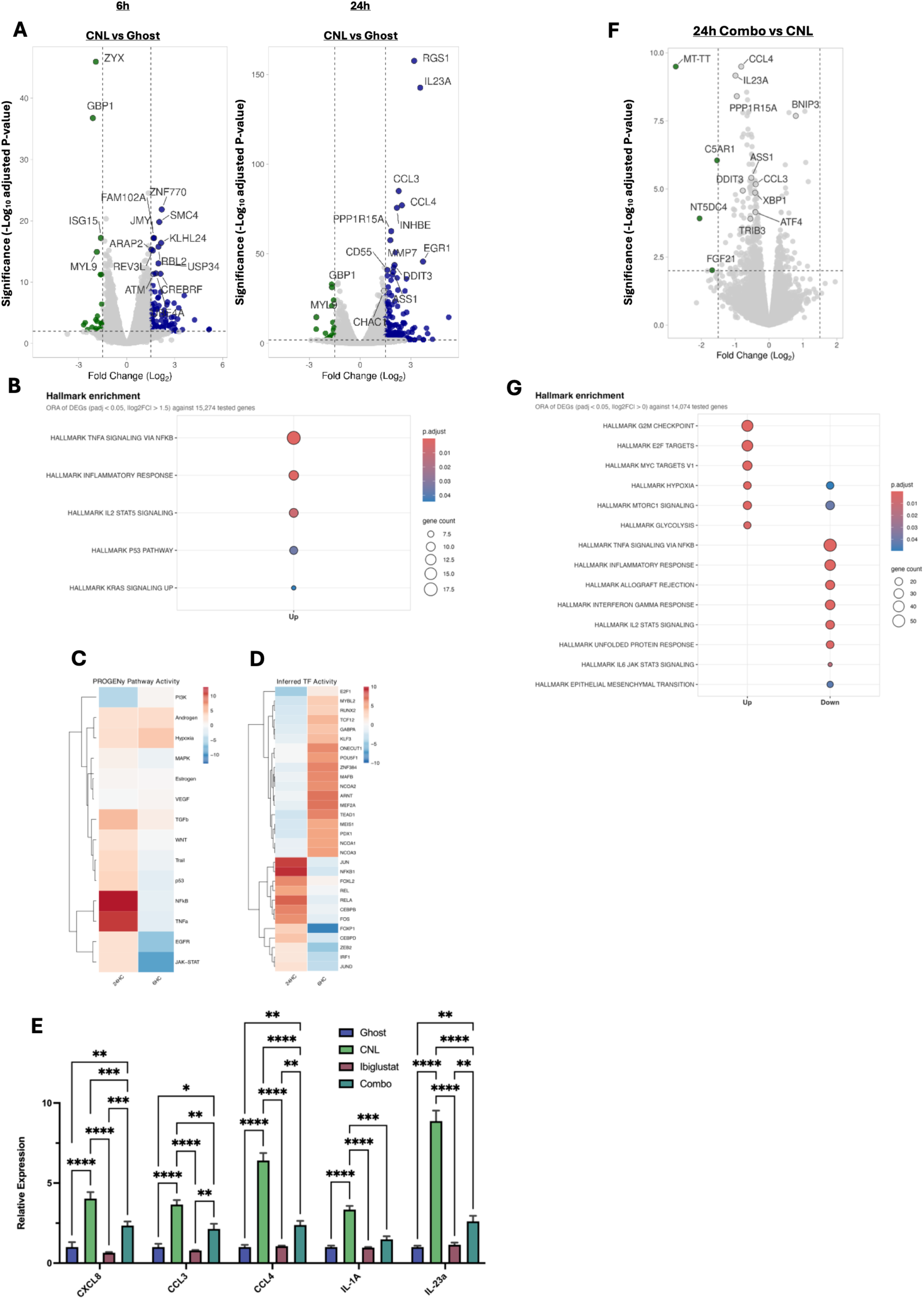
CNL treatment produces inflammatory and stress-related transcriptomic changes. **(A)** RNA-seq of JVM-3 cells showing key differentially regulated genes after 6 and 24 h of 20 µM CNL or control (ghost) liposome treatment. **(B)** Hallmark over-representation analysis of genes upregulated in the 24 h CNL group. **(C, D)** PROGENy pathway activity (**C**) and inferred transcription factor activity (**D**) for the pathways and factors most differentially active between the 6 h and 24 h timepoints. **(E)** RT-qPCR confirmation of cytokine expression after 24 h of 20 µM CNL, 2 µM ibiglustat or the combination. **(F, G)** Differentially expressed genes **(F)** and Hallmark pathway analysis (**G**) of RNA-sequencing data comparing 24 h of 20 µM CNL with 20 µM CNL plus 2 µM ibiglustat co-treatment; key stress-response genes are highlighted in **(F)**. Volcano plot thresholds (dotted lines) are |log2 fold change| > 1.5 and adjusted p < 0.05; upregulated genes are shown in blue and downregulated genes in green. RNA-seq data represent 3 (A–D) or 2 (F, G) biological replicates. Statistics for (E): two-way ANOVA with Fisher’s LSD test; error bars represent standard deviation of 3 independent experiments. Benjamini-Hochberg correction was used for RNA-seq. ****p<0.0001, ***p<0.001, **p<0.01, *p<0.05.

We next asked which transcriptional changes are reversed when GSL accumulation is blocked. A second RNA-seq compared CNL treatment with CNL/ibiglustat co-treatment. Cell-stress transcripts whose sustained expression is associated with death, including *DDIT3/CHOP* and *TRIB3*, were reduced in the combination (**Fig. 3F**). At the pathway level, the inflammatory signatures induced by CNL were among those most reduced by co-treatment, together with the unfolded protein response, while proliferative signatures including E2F and MYC targets increased (**Fig. 3G**), consistent with cells rescued from death resuming growth. The suppression of the unfolded protein response in particular directed us to organelle function, and to the ER, as a prospective node of disruption driving death.

### GSL accumulation drives ER stress, lysosomal deacidification and loss of mitochondrial respiration

Transcriptional signatures indicate that a program has been engaged but not that it has been executed, and the unfolded protein response is defined at the level of protein and RNA processing rather than transcript abundance. We therefore tested the ER stress signature directly. CNL induced TRIB3 accumulation and XBP1 splicing and both were suppressed when GSL synthesis was inhibited **(Fig. 4A)**. To resolve the timing, XBP1 splicing was measured by targeted RT-qPCR and was not induced at 5 h but was maximally induced by 12 h **(Fig. 4B)**, following GSL accumulation and preceding cell death. Cell surface calreticulin, a marker of ER stress and immunogenic cell death, increased only modestly **(Fig. 4C)**, and ER-Tracker Green staining intensity was modestly decreased after CNL **(Fig. 4D)**, consistent with UPR engagement alongside possible retraction of the compartment. Since ER-Tracker signal also reflects probe accessibility as well as ER membrane content, we do not interpret this shift as a change in ER mass. The ER response to CNL therefore appears to be a signaling response rather than structural collapse of the organelle.

**Figure 4:**
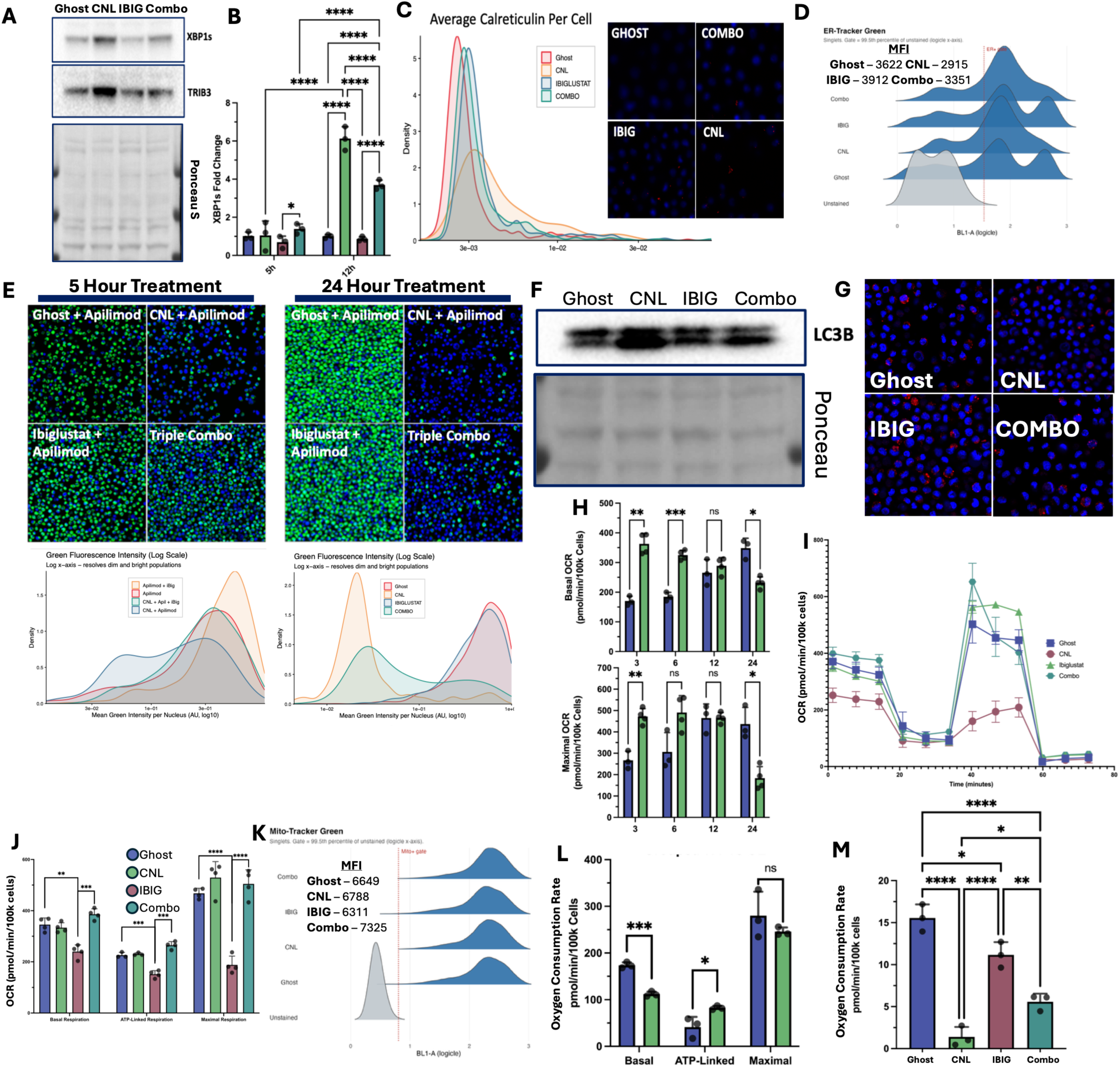
CNL-induced organelle stress is temporally ordered and reduced by ibiglustat. **(A)** Markers of ER stress, spliced XBP1 (XBP1s) and TRIB3, by western blot in JVM-3 cells after 24 h of 20 µM ghost liposomes, 20 µM CNL, 2 µM ibiglustat, or the combination. **(B)** Time course of XBP1 splicing by RT-qPCR (n=3). **(C)** Cell surface calreticulin levels in CNL-treated JVM-3 cells by confocal fluorescence microscopy; representative images and per-cell intensity distributions (n=1). **(D)** JVM-3 cells stained with ER-Tracker Green and analyzed by flow cytometry after 24 h of the above treatments; median fluorescence intensity from n=1 experiment. (E) LysoTracker Green staining of JVM-3 cells with or without 20 µM CNL and 2 µM ibiglustat at 5 and 24 h, with corresponding density plots. Apilimod (5 µM) was included in all conditions, including vehicle controls, to enlarge lysosomes and improve LysoTracker signal resolution; comparisons are therefore internally controlled but do not report absolute lysosomal pH (n=1). **(F)** LC3B-II accumulation by western blot after 24 h of 20 µM CNL with or without ibiglustat pre-treatment. **(G)** Cathepsin B activity assayed by Magic Red staining after 24 h of CNL; representative image (n = 1). Punctate versus diffuse signal was assessed qualitatively and was not quantified. **(H)** XFe96 Seahorse analysis of JVM-3 cells at the indicated times following 20 µM CNL or ghost treatment (n=4). **(I, J)** Seahorse analysis after 18 h of treatment ± 2 µM ibiglustat (**I**), with quantification **(J),** (n=4). **(K)** MitoTracker Green staining by flow cytometry to assay mitochondrial content after 18 h of the above treatments; median fluorescence intensity from n = 1 independent experiment. **(L)** XFe96 profiling of *UGCG*-overexpressing lines (green) compared with vector-control JVM-3 cells (blue) (n=3). **(M)** Basal respiration of primary CLL patient samples rescued with ibiglustat (3/7 patients, see Results) after 18 h of 40 µM ghost or CNL, ± 2 µM ibiglustat (n = 3 patient samples). Statistics: one-way ANOVA with Tukey’s post-hoc test (B, M), two-way ANOVA with Fisher’s LSD test (H), Student’s t-test (J, L). Error bars represent standard deviation. ****p<0.0001, ***p<0.001, **p<0.01, *p<0.05.

GSLs are synthesized in the Golgi and trafficked through the endo-lysosomal system, and their accumulation within lysosomes is the defining lesion of the sphingolipidoses. We therefore turned to that organelle. Using LysoTracker Green, which accumulates in acidic compartments, we found reduced signal within 5 h of CNL treatment, before the ER response, sustained to at least 24 h (**Fig 4E**). Both the early and late effects were alleviated by ibiglustat co-treatment. Apilimod was included in all conditions in this experiment to increase lysosome quantity and thus, signal intensity.

Consistent with loss of lysosomal acidity, LC3B-II, the lipidated, autophagosomal form of LC3B, accumulates in response to CNL and was likewise suppressed by ibiglustat (**Fig 4F**), suggesting impaired autophagic flux rather than increased autophagosome turnover. Co-treatment of CNL with hydroxychloroquine or apilimod produced no further accumulation of LC3B-II beyond either agent alone (**Fig. S2**), supporting a CNL-mediated flux effect.

To distinguish lysosomal deacidification from membrane permeabilization, intracellular cathepsin B activity was assessed by confocal microscopy. Lysosomal permeabilization releases cathepsin B into the cytosol and produces diffuse activity, whereas deacidification leaves it contained. Magic Red signal remained punctate under CNL treatment **(Fig. 4G)**, qualitatively consistent with lysosomes that lose acidity while remaining intact.

Neither the lysosomal nor ER measurements account for the loss of intracellular ATP observed after CNL treatment **(Fig. 1C)**. CLL cells depend heavily on oxidative phosphorylation for ATP production and mitochondrial function has been tightly linked to ER homeostasis. We therefore measured mitochondrial respiration by oxygen consumption rate over time, normalizing to equal numbers of viable cells so that the measurement reflects respiratory capacity per surviving cell rather than the loss of cells. The response was biphasic (**Fig. 4H**), where basal and maximal respiration were significantly increased relative to vehicle between 1 and 6 h, were indistinguishable at 12 h, and were significantly decreased by 18 and 24 h. The early increase coincides with the period of ceramide accumulation and precedes any detectable hexosylceramide elevation (**Fig. 1E, 1I**), whereas the later decline coincides with GSL accumulation. Consistent with this, Seahorse analysis at 18 h showed marked decreases in basal, ATP-linked and maximal respiration that were each rescued by ibiglustat co-treatment (**Fig. 4I**, quantified in **4J**). Mitochondrial content, assessed by MitoTracker Green, was unchanged across conditions (**Fig 4K**); this, together with the LC3B-II data, argues against mitochondrial clearance and for functional impairment of an intact mitochondrial pool. Additionally, cellular ROS was increased by CNL and trended downward, though not significantly, with ibiglustat (**Fig. S3**).

To determine whether increased flux through this pathway is sufficient to impair respiration, we measured oxygen consumption in JVM-3 cells overexpressing *UGCG* and found reduced basal respiration relative to vector controls **(Fig. 4L).** Reduced respiration in these cells was nonetheless not accompanied by increased sensitivity to CNL (**Fig. 2L**), indicating that diminished respiratory capacity alone is not sufficient to cause death. Reduced basal respiration after CNL was also observed in all seven primary CLL patient samples tested and was rescued by ibiglustat in three of seven (**Fig. 4M**).

These measurements also separate early from late organelle stress phases. Within the first hours, while ceramide is maximal and GSLs have not yet accumulated, respiration is elevated and ATP is already declining (**Fig. 1C**), a response consistent with inefficient or uncoupled respiration rather than with respiratory failure. Lysosomal deacidification is detectable by 5 h, ER stress is absent at 5 h and maximal at 12 h, and respiration falls below vehicle only by 18 h. The organelle lesions that depend on GSL accumulation therefore emerge after, and are separable from, the immediate mitochondrial response to delivered ceramide.

### GSL accumulation activates JNK signaling, which contributes to CNL-induced death

Organelle stress must be converted into a death signal. The stress-activated MAPK, JNK is the canonical route by which sustained, unresolved UPR signaling is transduced to death^20,21^ and UPR signaling through JNK and NF-κB also drives inflammatory gene expression of the kind observed here (**Fig. 3D**).^22^

A targeted analysis of stress-pathway transcriptional targets in our RNA-seq analysis showed consistent induction of JNK targets, whereas targets of the related p38 pathway responded in a mixed manner **(Fig. 5A)**. We therefore assayed two stress readouts over time: phosphorylation of c-Jun, the canonical JNK substrate, and induction of CHOP **(Fig. 5B)**. c-Jun phosphorylation was induced early and was suppressed by ibiglustat co-treatment, placing JNK pathway activity downstream of GSL accumulation. CHOP was induced maximally at 12 h, coincident with XBP1 splicing, was sustained to at least 24 h, and was likewise suppressed by ibiglustat, further linking GSL accumulation to sustained stress signaling.

**Figure 5:**
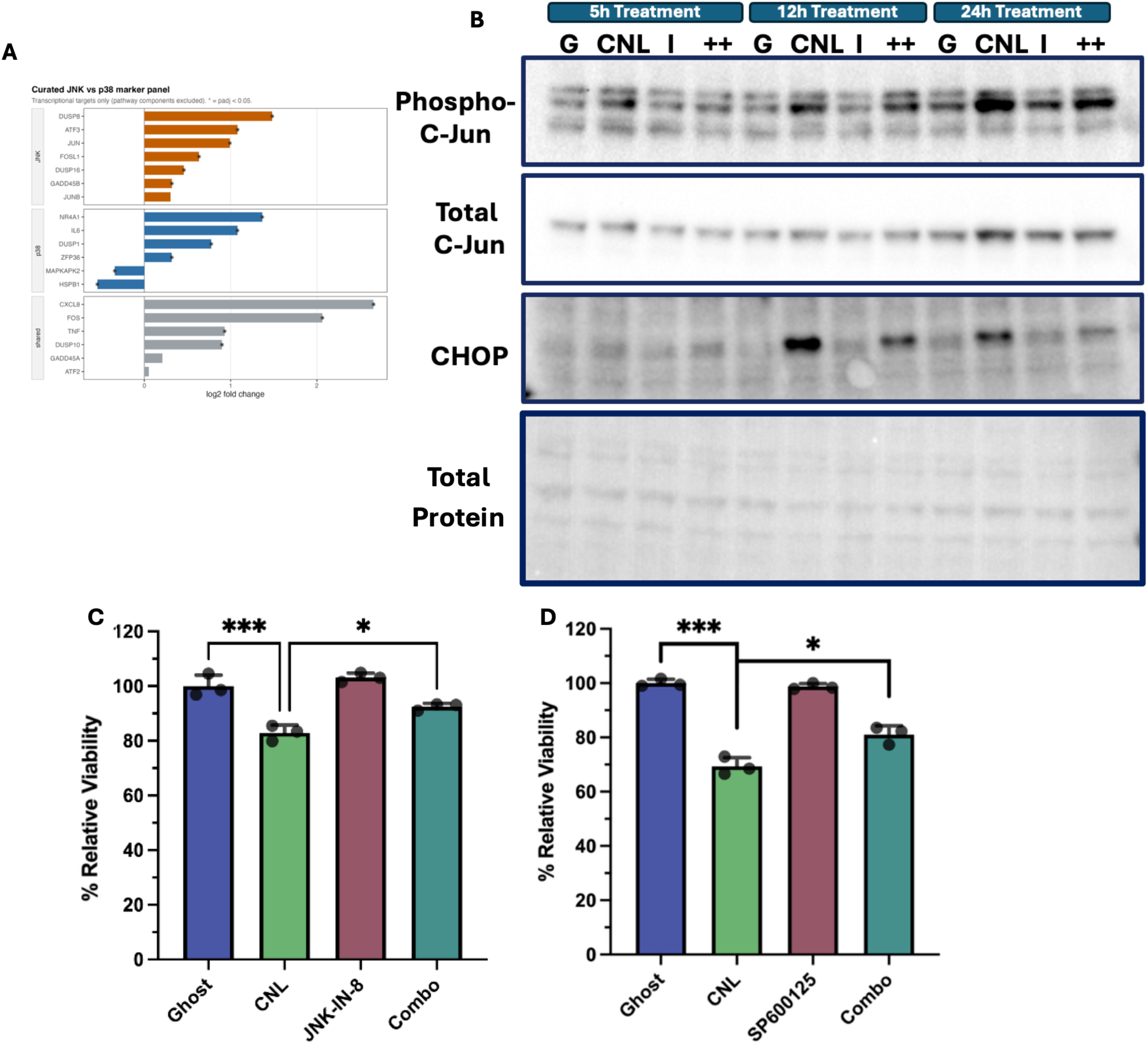
JNK signaling is engaged downstream of GSL accumulation and partially contributes to CNL-induced death. **(A)** Analysis of transcriptional targets of the JNK and p38 stress arms in RNA-seq data from JVM-3 cells treated for 24 h with 20 µM CNL versus ghost liposomes (average of 3 biological replicates); pathway components are excluded and only transcriptional targets are shown. **(B)** Western blot analysis of phospho-c-Jun, total c-Jun and CHOP in JVM-3 cells treated with 20 µM ghost liposomes, 20 µM CNL, 2 µM ibiglustat, or the combination, for the indicated times. **(C, D)** Viability of JVM-3 cells treated with 20 µM CNL in the presence or absence of 0.5 µM JNK-IN-8 **(C)** or 1 µM SP600125 **(D),** (n=3), assayed by alamarBlue. Statistics: one-way ANOVA with Tukey’s post-hoc test. Error bars represent standard deviation of 3 biological replicates. ***p<0.001, *p<0.05. Visual contrast was adjusted uniformly across the whole image in (B) to improve clarity; no region-specific adjustment was applied.

Lastly, we used two structurally unrelated JNK inhibitors to ask if JNK activity is required for CNL-induced death. Both the selective covalent inhibitor JNK-IN-8 (0.5µM) and SP600125 (1µM) partially protected cells from CNL (**Fig 5C and 5D**). The protection was incomplete with both inhibitors, indicating that JNK contributes to, rather than accounts for, the death program engaged by GSL accumulation. Contributions of accessory stress arms, including p38 and the integrated stress response, remain to be tested. As p38 also signals through post-translational mechanisms that a transcriptional signature would not capture, a contribution from that mechanism is not excluded.

### GSL synthesis is required for CNL efficacy across diverse lineages, and glucosylceramide alone is sufficient induce death

Having found that GSL accumulation is lethal rather than protective in CLL, we next determined whether these findings extend to other lineages. Ibiglustat rescued cells from CNL in MCF7 breast carcinoma (**Fig. 6A**) and G34 glioblastoma stem-like cells (**Fig. 6B**), with dose and duration adjusted for the sensitivity of each line. As our *UGCG* knockdown in CLL lines was incomplete, we extended the genetic test to *UGCG* knockout HEK293T and A549 cells. CNL reduced viability of wild-type A549 cells by ∼38%, an effect abolished in the knockout **(Fig. 6D)**. Wild-type HEK293T cells were considerably less sensitive to CNL, consistent with the reduced sensitivity of non-transformed lines generally, but the effect present in wild-type cells was likewise absent in the knockout **(Fig. 6C)**. Loss of *UGCG* therefore confers protection across a series of pharmacologic and genetic perturbations in five lineages.

**Figure 6:**
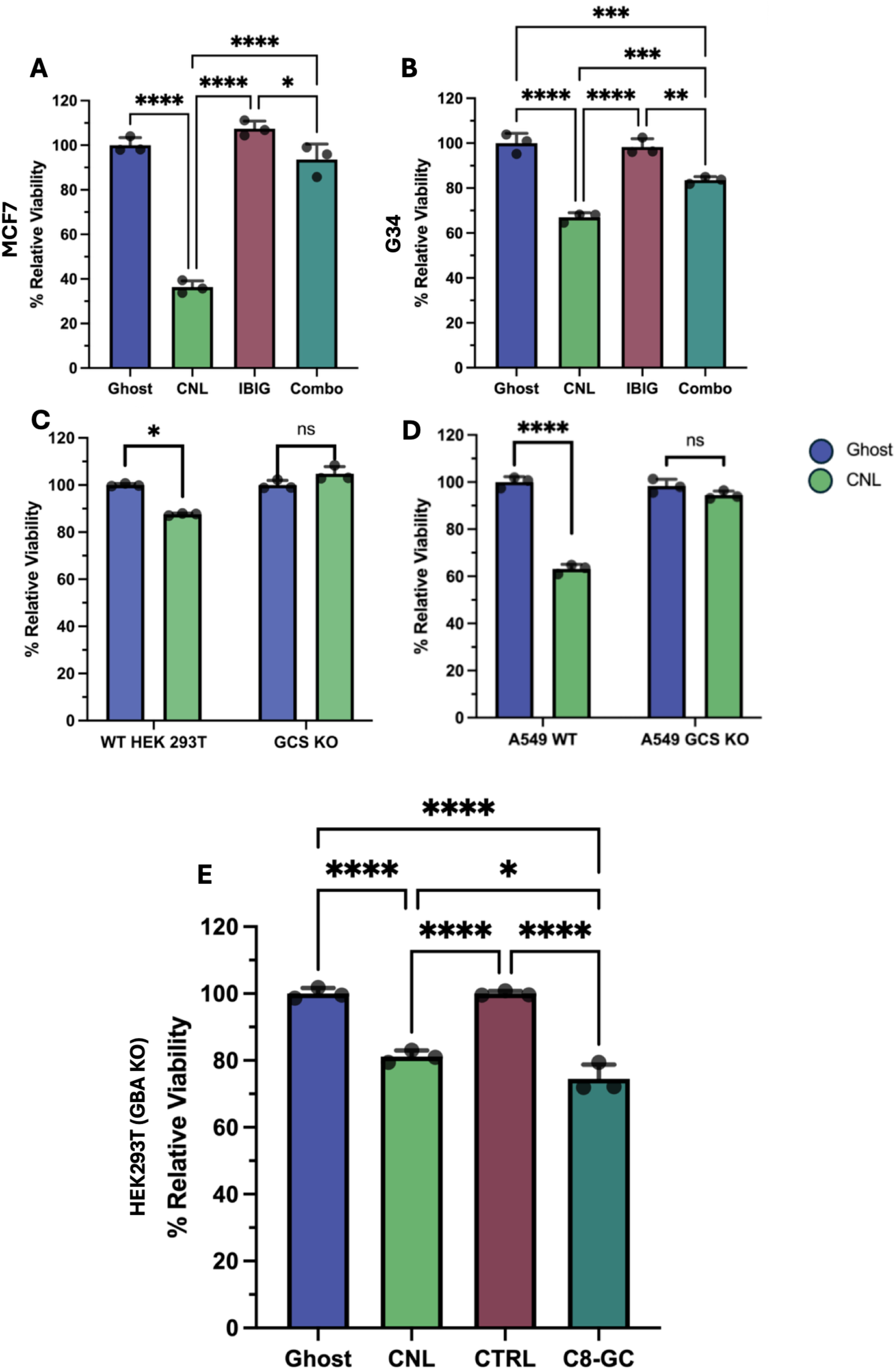
GSL synthesis is required for CNL-induced death across lineages, and glucosylceramide is sufficient to cause death. **(A, B)** alamarBlue viability following 72 h of 40 µM ghost or CNL treatment ± 2 µM ibiglustat in MCF7 breast carcinoma cells **(A)**, or 24 h of the same treatment in G34 glioblastoma stem-like cells **(B)**. **(C, D)** Sensitivity to CNL in wild-type and *UGCG*-knockout HEK293T (C) and A549 **(D)** cells, following 24 h or 48 h of 20 µM CNL (green) or ghost liposomes (blue), respectively. **(E)** *GBA1*-knockout HEK293T cells treated for 24 h with 40 µM ghost liposomes, 40 µM CNL or C8-glucosylceramide (C8-GC) or vehicle control (CTRL); viability assayed by alamarBlue. Statistics: one-way ANOVA with Tukey’s post-hoc test (A, B, E); Student’s t-test within genotype (C, D). In (C) and (D) each genotype was normalized to its own vehicle control, so these comparisons are within-genotype; the knockout result is an absence of a detectable CNL effect rather than a statistically demonstrated difference between genotypes. Error bars represent standard deviation of 3 biological replicates. ***p<0.001, ****p<0.0001.

These experiments establish that GSL synthesis is necessary for CNL-induced death, but not that a GSL is sufficient to induce death alone. To test sufficiency, and to ask whether death requires reconversion of the GSL back to ceramide, we used *GBA1*-knockout HEK293T cells, which cannot hydrolyze glucosylceramide. In these cells, C8-glucosylceramide reduced viability to an extent indistinguishable from CNL (**Fig. 6E**). A glucosylceramide delivered directly is therefore sufficient to cause death indicating that the lethal signal lies at or downstream of glucosylceramide rather than in regenerated ceramide.

## Discussion

Ceramide-based therapeutics are generally understood to cause cell death through ceramide accumulation, with metabolism viewed primarily as a route of detoxification that limits efficacy. Our data reveal the opposite: delivered C6-ceramide is actively metabolized into GSLs **(Fig. 1E–M)**, as expected,^23,24^ but this metabolic remodeling accompanies and is required for CNL-induced death. Pharmacological or genetic blockade of glucosylceramide synthase reduced rather than enhanced death across distinct cellular backgrounds **(Fig. 2F-H and 6A-D)**. GSL synthesis therefore represents not a route of ceramide disposal, but an essential step through which CNL engages the death program.

A requirement for GSL synthesis in causing cell death runs counter to a large literature in which glucosylceramide synthase inhibition sensitizes tumor cells to chemotherapy, however, that literature is less uniform than it first appears. The strongest results are genetic where UGCG expression confers, and its loss reverses, resistance to doxorubicin,^13,14^ but the effect was later attributed to regulation of MDR1 expression and drug efflux rather than to ceramide signaling.^25,26^ The pharmacological approach carries a separate problem as these inhibitors’ activity is not confined to the synthase,^27–30^ and the correlation between glucosylceramide content and resistance is itself confounded.^31^ When more selective inhibitors were used, chemotherapy sensitization was not reproduced.^32^ Our own pharmacology points the same way, where ibiglustat protected while raising ceramide levels above the levels reached with CNL alone, while sphingosine also increased and sphingomyelin, depleted by CNL, was restored toward baseline **(Fig. S1A–C)**. Ceramide and death move in opposite directions, and a detoxification model, in which inhibiting glucosylceramide synthase kills by increasing the ceramide pool, does not operate in this setting.

The MCF7 result is notable in this context **(Fig. 6A)** because this line is where glucosylceramide synthase overexpression was first shown to confer chemoresistance,^13^ and is the origin of the view that ceramide glycosylation is a detoxifying step; here, inhibiting that step protects rather than sensitizes. That study also reported resistance to exogenous ceramide in GCS-overexpressing MCF7 cells, which we do not observe as increased sensitivity on UGCG overexpression **(Fig. 2L)**. The two sets of observations differ in the form in which ceramide is presented and in whether synthase capacity is limiting, and together they argue that the consequences of ceramide glycosylation depend on the route of delivery rather than on glycosylation being intrinsically protective.

Inhibition of UGCG both raises ceramide in CNL-treated cells and shifts delivered ceramide toward the sphingosine/sphingosine-1-phosphate axis; genetic loss of *UGCG* produces a similar lipidomic shift.^33^ Neither manipulation distinguishes whether loss of a UGCG-dependent GSL signal promotes resistance or whether accumulation of another sphingolipid species contributes a pro-survival signal. Our findings also differ from a recent report in which glucosylceramide synthase inhibition sensitized H3K27M diffuse midline glioma to irradiation.^34^ That study tested both miglustat, a broad iminosugar, and eliglustat, a selective ceramide-analog inhibitor of the same class as ibiglustat, and reported ceramide accumulation with each inhibitor alone. In our system ibiglustat alone did not raise basal ceramide **(Fig. S1A)** and ceramide accumulated only when UGCG inhibition was combined with CNL. The difference between the two studies may therefore be in the insult rather than in the inhibitor. Where death is driven by an independent genotoxic stress, ceramide accumulation can sensitize; where death is driven by delivered ceramide itself, the requirement is for onward flux into GSLs, and the ceramide that backs up behind a GCS block is not itself sufficient to kill.

GSL accumulation was accompanied by an ordered organelle response. Lysosomal acidification was lost first, by 5 h **(Fig. 4E)**, and the ER stress response followed at 12 h, with XBP1 splicing and CHOP induction appearing on the same schedule **(Fig. 4A-C)**; both were prevented by ibiglustat, placing them downstream of GSL synthesis. The mitochondrial response was biphasic **(Fig. 4H–K)**. Respiration was elevated over the first hours, while ceramide was maximal and GSLs had not yet accumulated, and fell below vehicle only by 18 h, in parallel with GSL accumulation and reversibly on ibiglustat co-treatment. The early and late phases are therefore separable: the initial bioenergetic change tracks delivered ceramide, whereas the sustained respiratory defect, like the lysosomal and ER lesions, depends on metabolism into GSLs. These compartments are affected in GSL storage disease, and each limb has an independent precedent, although we are not aware of a single model in which their relative timing has been established. Glucosylceramide raises endolysosomal pH in Gaucher models and perturbs targeting along the endocytic pathway;^35,36^ GM1 accumulation activates the unfolded protein response and is sufficient to cause neuronal death in GM1-gangliosidosis;^37^ and downstream consequences in Gaucher models include impaired autophagy^38^ and mitochondrial dysfunction with defective quality control,^39^ which are partially corrected by enzyme replacement in patient cells^40^ and are viewed as consequences of substrate accumulation.^16,17^ For the mitochondrial arm there is in addition a specific candidate interface: mitochondria–lysosome membrane contact sites regulate mitochondrial dynamics and mediate Ca²⁺ transfer through lysosomal TRPML1,^41,42^ and these contacts are dysregulated in *GBA1*-mutant neurons, where the defect tracks with loss of glucocerebrosidase activity and is rescued by restoring it.^43^ Whether glucosylceramide accumulation is the proximate cause in that setting was not tested.

Three qualifications constrain this interpretation. First, we show a sequence with a shared upstream dependence, not causal propagation: we have not established that lysosomal deacidification causes the ER response or either causes the respiratory defect. GM1 accumulation at mitochondria-associated ER membranes offers an alternative route linking ER stress to Ca²⁺-dependent mitochondrial apoptosis.^44^ Second, the storage-disease analogy depends on the accumulating species. C6-GSLs are too short for the lysosomal sorting pathway,^45^ making the analogy dependent on concurrent accumulation of endogenous long-chain hexosylceramides; because ibiglustat suppresses both, the causal pool remains unresolved. Glucosylceramide depletion itself can also perturb endocytic targeting.^36^ Third, temporal ordering does not establish mechanism. Gangliosides can directly perturb mitochondria, induce ROS, and trigger permeability transition,^46^ consistent with our ROS data **(Fig. S3).** The preceding lysosomal phenotype argues against this route but does not exclude it. Direct ceramide effects on complex III are less consistent with our data because our phenotype requires downstream GSL synthesis.^47^

In each compartment the lesion was functional rather than destructive. Lysosomes lost acidity without evidence of integrity loss: acidotropic signal fell, while cathepsin activity remained punctate rather than diffuse **(Fig. 4E, 4G)**, which is the conventional distinction between deacidification and permeabilization.^48^ We note that this distinction rests on a single readout whose cleavage is itself pH-sensitive, and a direct assay of lysosomal membrane integrity would be required to establish it definitively. The ER showed a canonical unfolded protein response without changes in ER content **(Fig. 4D)**, while mitochondria lost respiratory capacity without loss of mitochondrial mass **(Fig. 4H and J)**, arguing against organelle clearance as the cause. The lysosomal phenotype also precedes, rather than results from, MLKL-dependent membrane rupture. A recent study reported MLKL translocation and polymerization at lysosomes, causing permeabilization and cathepsin release, with cathepsin B inhibition conferring protection.^49^ Here, lysosomal deacidification was evident by 5 h, preceding death, arguing against MLKL-driven rupture as its cause and placing the lysosomal lesion upstream of execution.

CNL also induced a substantial inflammatory transcriptional program pointing to TNFα/NFκB and inflammatory response signatures at 24 h, with induction of CCL3, CCL4, IL23A and related transcripts **(Fig. 3A–E)**. This program is GSL-dependent: each transcript was reduced by ibiglustat, and the corresponding pathway signatures were among those most reduced when GSL synthesis was blocked **(Fig. 3E, 3G)**. Whether it contributes to death or accompanies it remains unresolved. In B cells these pathways are more often associated with survival than with death, and inflammatory signaling induced by a dying population may be a consequence of stress rather than its cause. We note that the cytokines induced here are of potential interest independently of their contribution to death, since a cell death that recruits immune effectors would have different therapeutic implications in an immune-intact setting than one that does not.

Loss of UGCG prevented the phenotype, whereas constitutive overexpression did not enhance it **(Fig. 2I-L)**. This may indicate that UGCG is necessary but not rate-limiting, with downstream GSL trafficking or glycosylation constraining flux. Glucosylceramide must be selectively translocated from the cytosolic to the luminal Golgi leaflet for further glycosylation, through pathways involving FAPP2 and vesicular transport.^50–52^ Alternatively, constitutive overexpression may have permitted metabolic adaptation.^53^ Since ectopic UGCG expression can increase downstream GSLs and confer drug resistance,^13,54^ these results do not distinguish transfer limitation from adaptation. Acute, inducible UGCG expression with comprehensive GSL profiling would resolve this distinction.

The requirement for GSL synthesis is matched by evidence of sufficiency. In GBA1-knockout cells, which cannot hydrolyze glucosylceramide back to ceramide, C8-glucosylceramide delivered directly reduced viability to an extent indistinguishable from CNL **(Fig. 6E)**. The lethal signal therefore lies at or downstream of the GSL rather than in regenerated ceramide, which is the converse of the prediction made by the detoxification model.

### Limitations of the study

Three gaps constrain what we can conclude.

First, we have not identified what converts GSL accumulation into death. CNL increased MLKL phosphorylation in a GSL-dependent manner **(Fig. 2M–N)** without detectable caspase-3 cleavage, but we did not test whether MLKL is required, nor characterize RIPK3 expression in these cells, and CLL is a lineage in which RIPK3 is frequently downregulated. We therefore describe the phosphorylation without assigning a death modality. Relatedly, we have not established whether increased flux toward sphingosine and sphingosine-1-phosphate under UGCG inhibition contributes to protection; because S1P signaling is generally pro-survival, this remains an alternative to loss of a GSL-dependent death signal.

Second, we do not identify a specific GSL. Resolving this would require systematic genetic manipulation of the eight enzymes downstream of UGCG, which cellular adaptation may confound. The spatial dimension is equally unresolved. Sphingolipid localization is as tightly controlled as sphingolipid synthesis, and while we demonstrate increased cellular levels of many species, where those species accumulate is unknown. Given the effects across three organelles, localization would be needed to distinguish direct from indirect impingement, and we are developing compartment-resolved lipidomics toward this end.

Third, our model systems constrain generalization. JVM-3, our principal cell line, is derived from B-prolymphocytic leukemia rather than CLL, and the CLL claim rests most directly on the primary patient samples. *In vivo* consequences remain untested; given the inflammatory transcriptional program described here, the immune-intact setting is where these findings would matter most, but the predominant CLL mouse model (TCL-1) fails to recapitulate several core aspects of the disease, including activation of drivers that may themselves alter sphingolipid metabolism.

## Materials and Methods

### Compounds, reagents and antibodies

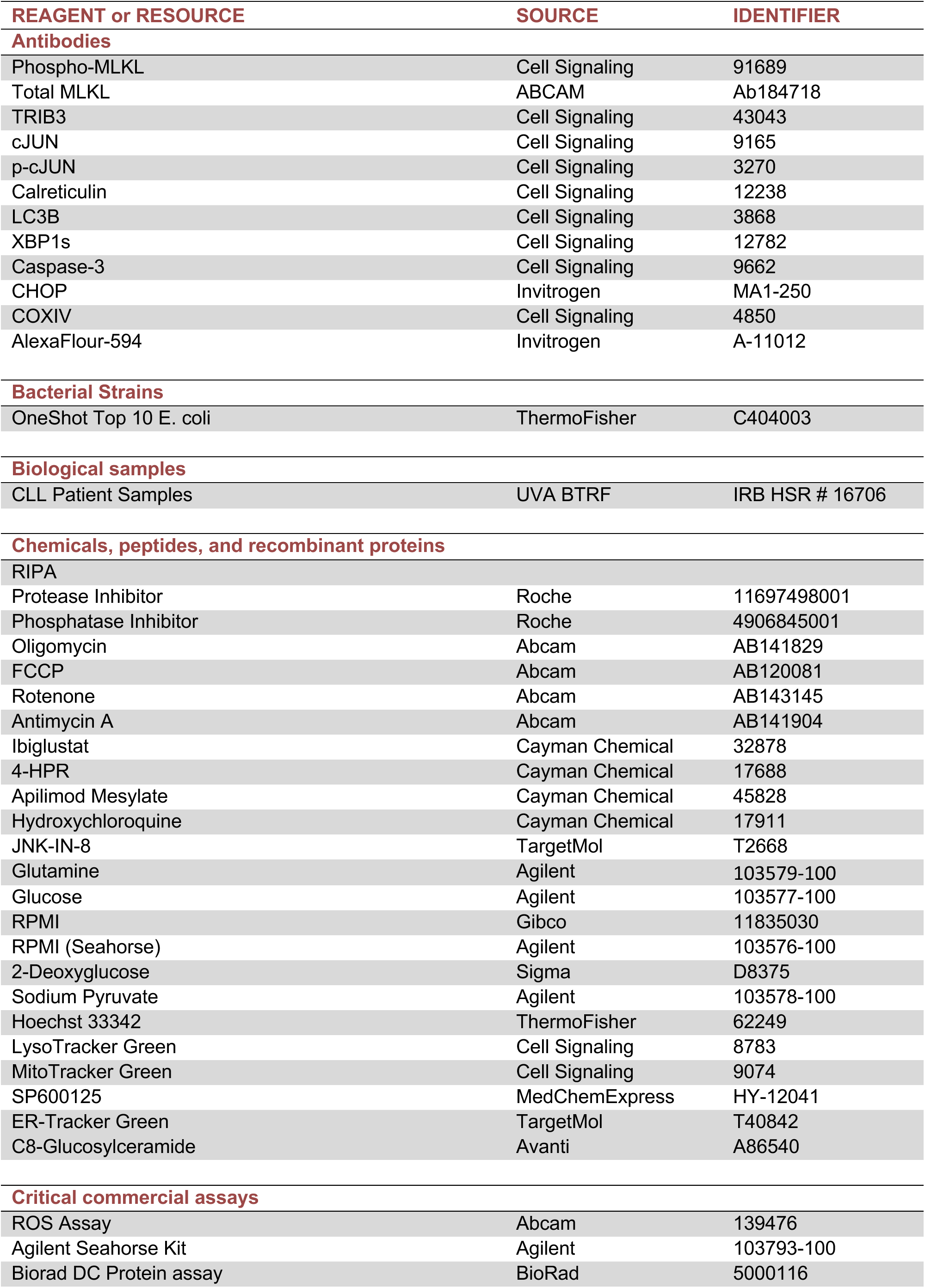
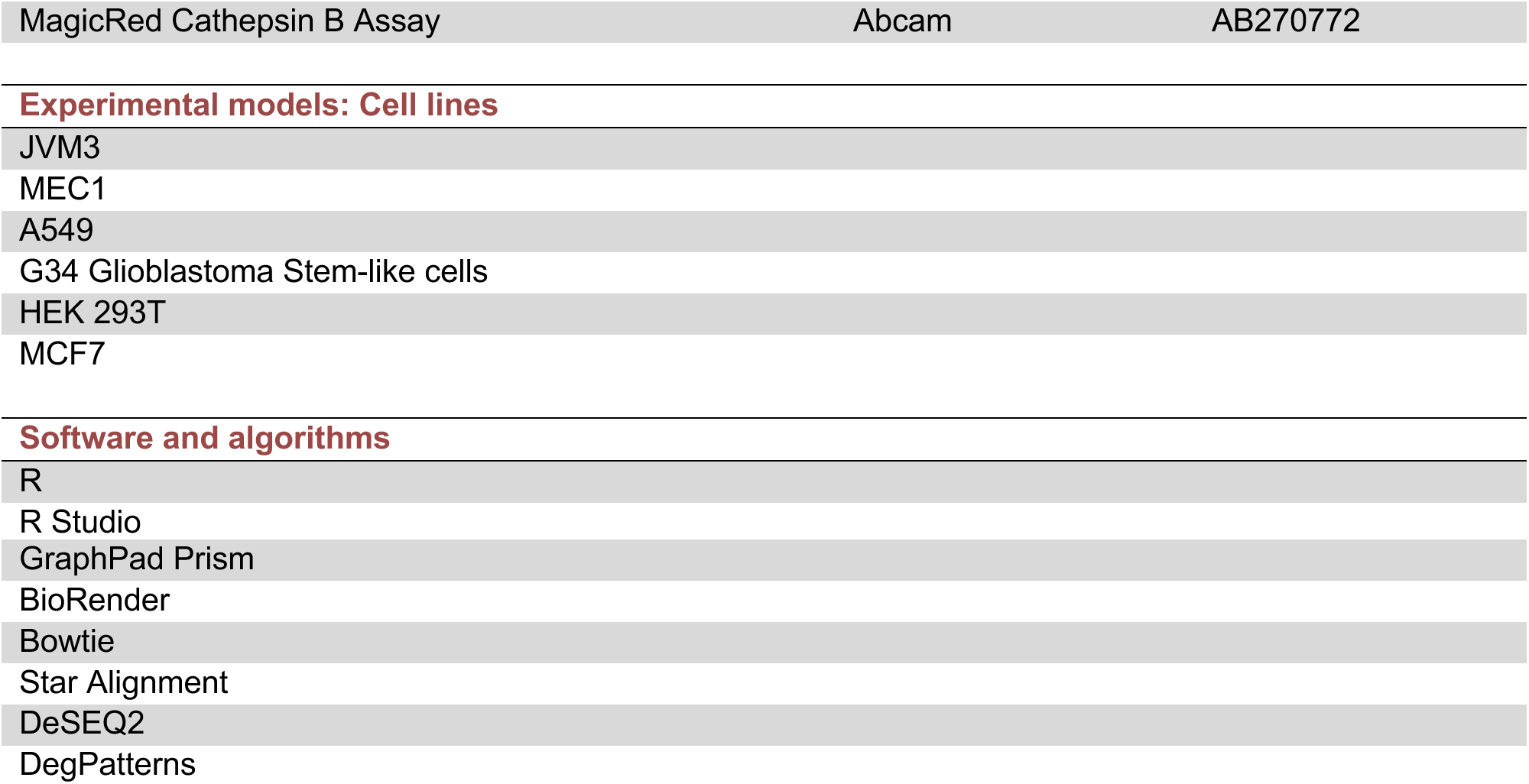

### Lipid Extraction and Analysis

Cell pellets were resuspended in 10% PBS to induce hypotonic swelling followed by brief probe sonication for 6 seconds at 35% amplitude. Each sample was then subjected to protein quantification using the BioRad DC protein assay (described below) and 200µg of sample was combined with 2mL of isopropanol:ethyl acetate:water (1:1:3) and 20pmol of Avanti sphingolipid internal standards. These tubes were then vortexed and sonicated 3x then rocked at 37°C for 1 hour. Tubes were spun at 3000xg for 10 minutes and the supernatant was transferred to a new tube. 2mL of Isopropanol:ethyl acetate:water was added to the pellet, vortexed, sonicated and allowed to rock at 37°C for another 15 minutes. These tubes were then centrifuged as before, and the supernatant was combined with the original 2mLs of extract. The combined liquid layers were dried down under constant nitrogen addition and resuspended in 60% methanol solution and filtered into an HPLC-MS/MS compatible plate. LC-ESI-MS/MS was used to analyze extracted samples. MS peaks were integrated, normalized to internal standards and protein quantification from the BioRad DC assay to yield the amount of each lipid species in pmol/mg protein.

### Western Blot

Cell pellets were incubated for 15 minutes on ice with Thermofisher Scientific premade RIPA buffer that had been supplemented with ROCHE cOmplete protease and PhosStop phosphatase inhibitor tablets. These samples were then sonicated for 12 seconds at 35% amplitude followed by centrifugation for 12,000xg for 10 minutes at 4°C. Protein concentration of each sample was determined using DC protein assay and 30µg of sample was combined with 4x Laemmli buffer containing 2-mercaptoethanol and RIPA to a total volume of 50µL in Eppendorf tubes followed by heating at 70°C for 10 minutes. Samples were then loaded onto Thermofisher 4-12 Bis-Tris SDS-PAGE gels and ran for 35 minutes at 180V in Thermofisher MOPS buffer. Gels were then transferred onto 0.2µm PVDF membranes using BioRad’s TurboTransfer system and blocked with 5% BSA in TBST for 1 hour at room temperature. Primary antibodies were diluted in 5% BSA in TBST, added and allowed to incubate overnight followed by 3 washes with TBST and incubation with secondary antibody for 1 hour at room temperature. Blots were then washed 2x with TBST then 1x with TBS. Prior to imaging, blots were soaked with Thermo Fisher SuperSignal Femto ECL Reagent ECL for 1 minute and imaged using BioRad’s ChemiDoc system. Blots were then incubated with ponceau S for 1 hour, washed with water 3x and imaged to determine total protein per lane. Licor Image Studio Lite was used to quantify bands via densitometry.

### Cell Culture

a) Cell lines: JVM-3, A549, MEC1, MCF7 and HEK293T cells were all grown in RPMI supplemented with 10% FBS. Glioblastoma stem-cell like G34 cells were obtained from Dr. Benjamin Purow (UVA) and grown in neurobasal media supplemented with Gibco B-27 and N-2 supplements along with 25ng/mL EGF, 25ng/mL FGF and 1x cell culture grade glutathione. All cells were grown at 37C, 5% CO_2_.
b) Patient samples. Peripheral blood mononuclear cells were isolated from patients with chronic lymphocytic leukemia by density gradient centrifugation using Ficoll-Paque and cryopreserved. Samples were obtained with written informed consent under University of Virginia IRB-HSR protocol 16706 in accordance with the Declaration of Helsinki. Samples were thawed and rested for 1 h in RPMI with 10% heat-inactivated FBS before treatments.

### RT-qPCR

RNA was purified from cells using trizol mediated phase separation. In brief, cell pellets were resuspended in 1mL TRIzol and homogenized. After 5 minutes, 200µL of chloroform was added and tubes were vortexed vigorously for 2 minutes followed by centrifugation for 15 minutes at 17k x g at 4°C. An upper aqueous layer was removed and placed in a new tube to which 500μl of isopropanol was added. These tubes were inverted and placed in a centrifuge for 10 minutes at 17k x g at 4°C. The supernatant was then discarded and 1ml of ice cold 75% ethanol was added to each tube before centrifuging again for 5 minutes. The final pellet was dried and solubilized in nuclease free water. RNA was quantified using a plate based nanodrop and Cytation 3 software. cDNA was formed using a BioRad cDNA synthesis kit and diluted 1:10 following reverse transcription. 2µL of this cDNA was then combined with 0.5µL FAM conjugated probes, 5µL of BioRad iScript Probe Supermix, and 2.5µL water. Plates were placed in a BioRad RT-qPCR instrument, and the standard protocol (2 min at 95°C followed by 40 cycles of 95°C for 10 seconds and 60°C for 30 seconds) was used to determine expression. Probes were FAM conjugated and purchased from BioRad. XBP1s probes were synthesized by Eurofins (FWD: 5’ GCTGAGTCCGCAGCAGGT, REV: 5’ CTGGGTCCAAGTTGTCCAGAAT) and amplification was measured using SybrGreen supermix and the standard imaging protocol (2 min at 95°C followed by 40 cycles of 95°C for 10 seconds and 60°C for 30 seconds with an additional melt-curve of 65°C-95°C at 0.5°C/5 seconds). All graphs report 2^-ΔΔCt^ values for the indicated target relative to vehicle controls.

### Cell Viability

Cells were plated in 96- or 24-well plates at 500k cells/mL for suspension and 100k cell/mL for adherent lines and allowed to attach (overnight, adherent lines) or acclimate (1h for suspension cells) prior to treatment. Following the desired treatment duration, 10µL (100 µL for 24-well plates) of Alamar Blue was added per well and allowed to incubate for 2-4h after which fluorescence was read using 544 excitation and 590 emission filters. All values were normalized relative to control. For experiments utilizing 3-(4,5-dimenthylthiazol-2-yl)-5-(3-carboxymethoxyphenyl)-9-(4-sulfophenyl)-2H-tetrazolium (MTS), 150μL phenazine methosulfate (PMS, 0.92mg/mL stock) was combined with 1.5mL of MTS (2mg/mL stock) to generate a working stock of which 20μL was added per well, incubated for 2 hours after which absorbance was read at 490nm on a Cytation 3 plate reader.

### RNAseq

JVM3 cells were treated with 20 µM Control (ghost) liposomes or 20 µM CNL for 6h or 24h then collected, washed with PBS and flash frozen. Cell pellets were sent to Novogene for RNA sequencing using Illumina PE150 methodology. Novogene quality control (FASTQC package), Adapter trimming (CutAdapt Package) and read alignment to the GRCh38 human genome (STAR Alignment/BowTie packages) and provided us with raw read count files. The final read counts from this pipeline were analyzed using the R package DESeq2 which is an all-inclusive analysis package for calculating relative fold changes, significant hits (including correction for multiple statistical tests) and data visualization. Volcano plots were generated using the VolcanoseR webtool. Pathway analysis was conducted using the MSigDB Hallmark reference pathways and available R scripts to conduct over-representation analysis (ORA). To compare CNL treatment with CNL/ibiglustat co-treatment, JVM-3 cells were treated for 24 h with 20 µM CNL or with 20 µM CNL plus 2 µM ibiglustat (n = 2 biological replicates per arm), and RNA was purified using a Qiagen RNeasy kit. Purified RNA was sent to Plasmidsaurus LLC where it was sequenced and processed via their in-house pipeline. Pathway analysis using Hallmark reference pathways was conducted as above and Volcano plots were generated using VolcaNoseR. Pathway and transcription factor inference comparing 24h and 6h DEGs was done using PROGENy, DoRothEA and decoupleR packages in R.

### Seahorse Assay

Cells were treated as previously described. The day before seahorse analysis an Agilent XFe96 flux pack was hydrated using 200μL of calibrant per well and stored in a CO_2_ free incubator overnight. The day of seahorse analysis, treated cells were counted and a sufficient number of viable cells were pelleted such that once resuspended in 250μL of seahorse media (Agilent RPMI, 10mM glucose, 1mM sodium pyruvate and 2 mM glutamine) they would be at a concentration of 100k cells/50µL. Then 50µL of cells were added to 4 wells and spun down for 5 minutes at 300xg with no brake. Once settled, an additional 130μL of seahorse media was added and 180μL of seahorse media was added to blank wells throughout the plate. This cell plate was placed in a CO_2_ free incubator while the top Flux cartridge was loaded with oligomycin (1 μM final concentration in well), FCCP (1 μM final concentration) and rotenone/Antimycin A (0.5 μM final concentration). This sensor plate was then combined with the cell plate, and a standard Agilent mitochondrial stress test protocol was used for readings/injections on the seahorse instrument.

### Confocal Microscopy

Cells were treated as desired then stained immediately prior to imaging. For live cell dyes, 500k cells were pelleted, washed once with PBS then resuspended with the appropriate dyes (MitoTracker green 400nM, MitoTracker deep red 500nM, ER-tracker green 1µM, lysotracker green 50nM, Hoechst 10µM) in PBS and allowed to incubate for 10 minutes at room temperature. Cells were then pelleted, washed again with PBS and resuspended in 250µl of PBS and placed in the well of an Ibidi 8 chamber glass slide. 20x objective on a Zeiss 880 was utilized and images were collected using Zen Black software. Quantification was done in R studio (see GitHub. For antibody staining, cell pellets were resuspended in PBS then placed onto poly-l-lysine coated coverslips in the well of a 6-well plate. These plates were spun at 300xg for 5 minutes and the PBS was removed and replaced with 4% PFA. Cells were fixed for 25 minutes at room temperature followed by two washes of PBS then permeabilization in 0.25% triton x-100 in PBS for 5 minutes with gentle agitation if intracellular staining was required, this step was skipped for surface calreticulin staining. Cells were washed two more times with PBS then transferred to a humidified chamber with the cell side of the coverslip facing upward. Antibodies were diluted in 3% BSA in PBS and 160µL of this was added to the coverslip. Cells were incubated overnight at 4°C then washed 5x with PBS containing 0.1% Tween20. Coverslips were returned to the humidified chamber and secondary antibody solutions containing Hoechst as a counterstain were added to each coverslip and allowed to incubate for 1h at 37°C followed by 5 more washes in PBS-Tween20. 5 µL of mounting medium (50% glycerol) was added to a slide and coverslips were placed cell side down and edges were sealed using nail polish. Slides were imaged as using a Zeiss 880 confocal microscope as described above.

### Flow Cytometry

Cells were stained as described for live cell dyes in the confocal microscopy section then placed into a 96-well round bottom plate. These were loaded onto an Invitrogen Attune NxT for analysis and single stained/unstained controls were used for voltage acquisition. 30k events were captured. Gating and analysis were done using flowCore and ggcyto R packages by first selecting singlets using the diagonal portion of cells resulting from plotting Forward Scatter-Area vs Forward Scatter-Height. Next, single stain controls were used to set gates for each dye. Compensation was not required for live-cell dyes used.

### Intracellular ATP Estimation

Cells were stained with trypan blue following treatment and equally replated based on viable cell numbers into white 96-well plates. 100µL of CellTiter Glo was added to each well and incubated with shaking for 5 minutes to induce lysis. Luminescence was acquired using a Cytation 3 plate reader.

### Cell Line Generation

pLP-VSVG, pLP1, pLP2 and vector plasmids were combined with lipofectamine and p-3000 reagent as per the manufacturer’s suggested values and added dropwise onto 90% confluent HEK293T cells. The media was changed the following day and collected at 48 and 72h. Lenti-x concentrator was added to each tube of viral media and allowed to incubate for 4 hours followed by centrifugation. The resulting pellet was resuspended in culture media and aliquoted for future use. When transducing our cell lines of interest, viral particles were added to the culture along with 7µg/ml of polybrene and cells were spun at 200xg for 45 minutes to spinoculate the culture. Cells were returned to the incubator, and fresh media was added the following day. Selection with puromycin and/or flow sorting was conducted 3 days following transduction. Cell lines were validated via RT-qPCR.

### Statistics

All statistical analysis was performed in GraphPad Prism 10 except for RNA-seq analysis which was conducted in R. Comparisons of two groups used unpaired two-tailed Student’s t-tests; comparisons of three or more groups used one-way or two-way ANOVA as stated in each figure legend. Where three or more comparisons were made within a group, post-hoc tests or multiple testing correction was used to reduce type I errors: Tukey’s post-hoc test (one-way ANOVA), Benjamini-Hochberg correction (RNA-seq) or Fisher’s LSD test (two-way ANOVA). The test used and the number of replicates are stated in each figure legend. All error bars represent standard deviation from the mean and graph points represent biological replicates (analysis of independently conducted experiments) as opposed to technical replicates (repeated measures of the same experimental sample). Asterisks represent significance of p-values: *p<0.05, **p<0.01, ***p<0.001, ****p<0.0001

## Data/Code Availability

All RNA-seq counts and R pipelines utilized in this study will be made publicly available via deposition onto the proper online repositories (GEO and GitHub, respectively) following peer-reviewed publication.

## Author Contributions

L.R.V. and T.E.F. conceived the study and designed the experiments. L.R.V. performed the majority of the experiments, analyzed the data and prepared the figures. E.W.S. performed western blots and analyzed data. J.M.C. contributed to computational analysis of RNA-seq data and interpretation. S.L.J. performed RT-qPCR experiments. A.S. performed flow cytometry experiments and analysis. P.C.-P. and J.J.P.S. contributed to the experimental design and interpretation of results. T.P.L. provided interpretation of the CLL data. T.E.F. supervised the study, and wrote the manuscript with L.R.V. All authors approved the final manuscript.

## Acknowledgements

We thank Drs. Craig Portell and Michael Williams for access to their CLL patient sample repository (IRB-HSR-16706), Dr. Timothy Bullock for use of his Seahorse XFe96 instrument, and Dr. Kallesh Jayappa for many insightful conversations and suggestions regarding patient sample handling. We also thank the UVA Advanced Microscopy Facility for use of their confocal microscope, the Experimental Pathology core for use of their flow cytometer, and Plasmidsaurus and Novogene for sequencing services.

This work was supported by an NIH/NCI Predoctoral to Postdoctoral Fellow Transition Award (F99/K00; 5F99CA294256 to L.R.V.); a pilot project award from the University of Virginia U54 Systems Analysis of Stress-adapted Cancer Organelles (SASCO) program (U54CA274499); the NIH/NCI Program Project Grant P01CA302570); and the University of Virginia Comprehensive Cancer Center Support Grant (P30CA044579).

Lastly, this work would not have been possible without the late Dr. Mark Kester, who originated the ceramide nanoliposome, initiated the line of investigation pursued here, and mentored L.R.V. in his early training.

---

**Supplemental Figure S1: Ibiglustat co-treatment leads to accumulation of multiple upstream sphingolipid species.** LC-MS/MS lipidomic analysis of JVM-3 cells after 24 h of 20 µM ghost or CNL treatment, with or without 2 µM ibiglustat (IBIG), for total endogenous ceramides **(A)**, sphingosine **(B)** and sphingomyelin **(C)**. Statistics: one-way ANOVA with Tukey’s post-hoc test. Error bars represent standard deviation of 4 biological replicates. *p<0.05, **p<0.01, ***p<0.001, ****p<0.0001.

**Supplemental Figure S2: CNL does not increase LC3B-II accumulation beyond apilimod or hydroxychloroquine alone.** Western blot analysis of LC3B in JVM-3 cell lysates after 24 h of treatment with 20 µM CNL, 20 µM ghost liposomes (G), 1 µM apilimod (A) and/or 5 µM hydroxychloroquine (HQ); D denotes vehicle (DMSO). Equal protein was loaded in each lane with total protein shown in (B) as verified by BioRad StainFree gel assay. Lanes marked H2 contain 5 µM H282, a STAT3 inhibitor carried over from an unrelated experiment run on the same membrane; these lanes are shown for completeness and are not discussed further.

**Supplemental Figure S3: CNL increases cellular ROS.** Reactive oxygen species were measured in JVM-3 cells using Abcam’s ab139476 after 24 h of 20 µM ghost or CNL treatment, with or without 2 µM ibiglustat (IBIG). Statistics: one-way ANOVA with Tukey’s post-hoc test. Error bars represent standard deviation of 5 biological replicates. **p<0.01, ***p<0.001.

