## Supplemental Figures for "Glucosylceramide synthase is required for C6-ceramide nanoliposome-induced organelle stress and cell death"

Supplemental Figure S1: Ibiglustat co-treatment leads to accumulation of multiple upstream sphingolipid species

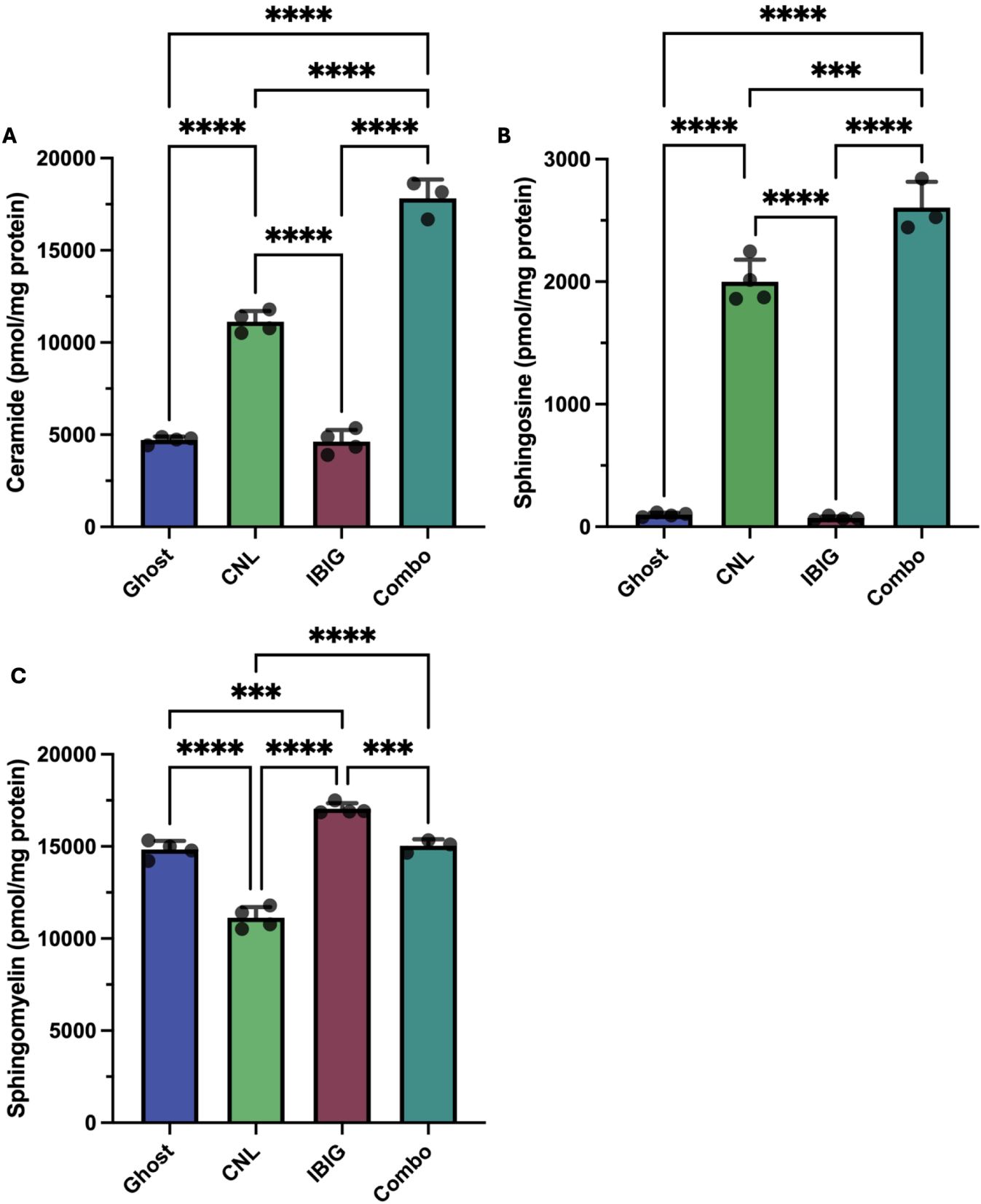

**Supplemental Figure S2: CNL does not increase LC3B-II accumulation beyond apilimod or hydroxychloroquine alone**

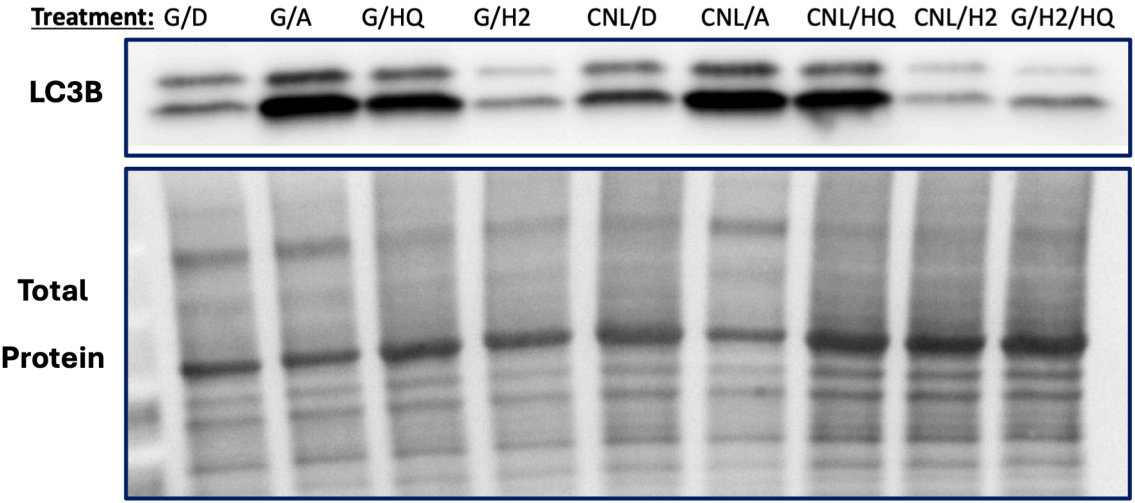

**Supplemental Figure S3: CNL increases cellular ROS**

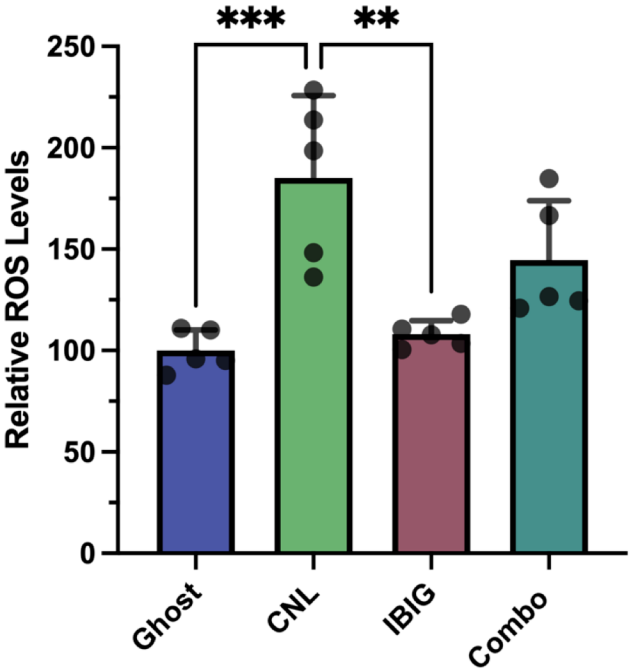
